# Binding Affinity Regression Models from Repeats Mutation in Polyglutamine Disease

**DOI:** 10.1101/281949

**Authors:** P R Asha, M S Vijaya

**Affiliations:** Department of Computer Science, Department of Computer Science, PSGR Krishnammal College for Women, PSGR Krishnammal College for Women, Coimbatore, India

**Keywords:** binding affinity, docking, ligand, polyglutamine repeats, prediction, numpy, scikit learn

## Abstract

Diagnosing and curing neurodegenarative disorder such as spinocerebellar ataxia is complicated when there is differences in formation of protein sequences and structures. Affinity prediction plays vital role to identify drugs for various genetic disorders. Spinocerebellar ataxia occurs but mainly it occurs due to polyglutamine repeats. This research work aims in predicting the affinity of spinocerebellar ataxia from the protein complexes by extracting the well-defined descriptors. Regression models are built to predict the affinity through machine learning techniques coded in python using the Scikit-Learn framework. Energy complexes and protein sequence descriptors are defined and extracted from the complex and sequences. Results show that the SVR is found to predict the affinity with high accuracy of 98% for spinocerebellar ataxia. This paper also deliberates the results of statistical learning carried out with the same set of complexes with various regression techniques.

## I. Introduction

Spinocerebellar ataxia also known as spinocerebellar atrophy or spinocerebellar degeneration is a progressive, degenertive disorder with multiple types. It is a hereditary anarchy portrayed by deviations in grey matter handling its tasks. The disorder is due to mutations in the genes which results in brain and spinal cord degeneration. Each type of SCA features its own symptoms [1]. Multiple forms of SCA caused by repeats mutation are SCA1, SCA2, SCA3, SCA6, SCA7, SCA8 and SCA10 [2]. When parent’s posses 39 repeat of polyglutamine and when it is passed to off-spring it may increase the repeats and causes mutation [3].

Trinucleotide repeat disorders conjointly referred to as trinucleotide repeat expanison disorders, triplet repeat enlargement disorders or sequence reduplication disorders. It is a collection of genetic disorders caused by trinucleotide repeat enlargement, a sort of mutation wherever trinucleotide repeats in bound genes or introns exceed the conventional, stable threshold, that differs per factor. The mutation may be a set of unstable microsatellite repeats that occur throughout all genomic sequences. If the repeat is gift in an exceedingly healthy factor, a dynamic mutation might increase the repeat count and lead to a defective factor. If the repeat is gift in associate degree deoxyribonucleic acid will cause toxic effects by forming spherical clusters known as polymer foci in cell nuclei. Trinucleotide repeats area unit typically classified as insertion mutations [4].

Repeats mutation causes many medical specialty disorders. Currently, 9 medical specialty disorders are familiar to be caused by associate multiplied variety of CAG repeats, usually in coding regions of otherwise unrelated proteins. Throughout super molecule synthesis, the enlarged CAG repeats are translated into a series of uninterrupted amino acid residues forming what’s referred to as a polyglutamine tract (“polyQ”). Such polyglutamine tracts could also be subject to multiplied aggregation [5].

In the Eukaryotic genes, repeat mutation occurs by three building blocks of DNA (cytosine, adenine, and guanine) that appear multiple times in a row. Due to trinucleotide repeat disorder there occur many neurological disorders, in which one of the disorders is spinocerebellar ataxia. Mode of inheritance for spinocerebellar ataxia is through autosomal, where disorder is passed by both parents. The most common form of SCA that are affected are affected by repeat mutation are SCA1, SCA2, SCA3, SCA6, SCA7, SCA8 and SCA10 [2], which posses many variance that code for protein.

Translation of protein from gene sequence is crucial form in human. Amino acid chains fold into 3D structures to form protein. Structures plays vital role in protein formation. Primary structure is a specific sequence made out of the 20 amino acids that build life. Secondary structure is a low-level structural element of a protein with alpha, helix and beta-sheet. Tertiary structure is the overall folding structure of the whole amino acid chain. Quaternary structure is overall fold of all related amino acid chains into a structure of more complex machine. After structure formation protein undergoes post-translational modification in which protein undergo some chemical processing by other proteins to become active [6].

Drug designing is a process in which by having knowledge of biological target a new medication is discovered. A drug is a small organic molecule that either activates or inhibits function of a biomolecule like protein which are involved in disease and hence result in therapeutic benefit to patient. The basic mechanism behind drug design is the formation or design of such molecules which have shape and charge complementary to biological target with which drug interact and bind with it and ultimately result in change in function of biological target. In drug designing process the technique in which use of computer is involved is called Computer Aided Drug Design. There are two types of Computer Aided Drug design i.e. ligand based drug design and structure based drug design [7].

Ligand based drug design involves the knowledge of molecule that bind to target molecule. Structure based drug design involves knowledge of 3D structure of biological target which is obtained from X-ray crystallography or NMR spectroscopy. If structure for target is available it is easy to create homology model against it. Using the structure of the biological target, candidate drugs that are predicted to bind with high affinity and selectivity to the target may be designed using interactive graphics and the intuition of a medicinal chemist [8].

In this paper structure-based drug design processes are used with machine learning techniques are used to predict affinity. Structure-based drug design can be divided roughly into three main categories. The first method is identification of new ligands for a given receptor by searching large databases of 3D structures of small molecules to find those fitting the binding pocket of the receptor using fast approximate docking programs. This method is known as virtual screening. A second class is De novo style of latest ligands. During this methodology, substance molecules area unit engineered up inside the constraints of the binding pocket by collection little items in a very stepwise manner. These items is either individual atoms or molecular fragments. The key advantage of such a way is that novel structures, not contained in any data, is usually recommended. a third technique is that the development of acknowledged ligands by evaluating planned analogs within the binding cavity [9].

Due to protein structure changes, there occurs a lot of changes in affinity binding. After structure formation when mutation is induced the protein structure and its function is altered [10]. In this paper the protein structures are altered by inducing the repeats mutation based on human genome mutation database, and the difference in affinity prediction. Repeats mutation information for the six types of spinocerebellar ataxia is depicted in Table 1.

**Table 1.** Repeats mutation information for SCA.

| Protein | No.of Repeats |
| --- | --- |
| Ataxin-1 | 40-100 |
| Ataxin-2 | 32-500 |
| Ataxin-3 | 68-79 |
| Ataxin-6 | 21-28 |
| Ataxin-7 | 40-200 |
| Ataxin-8 | 116 |
| Ataxin-10 | 1611 |

Docking is performed to forecast the binding modes. An earlier illumination for the ligand-receptor binding procedure is lock-and-key principle, wherein the ligand sits into the protein just like lock and key. After that induced-fit concept, it carries lock-and-key theory a phase more, proclaiming that the energetic site of the protein is constantly reshaped by interactions with the ligands since the ligands communicate with the macromolecule [11]. There is plenty of docking software’s available [12].

Affinity is a measure of the strength of attraction between co-ordination bonds with a receptor is known as binding affinity. The binding affinity of a ligand with a receptor depends upon the interaction force of attraction between the ligands and their receptor binding sites. Binding affinity of substance to receptor is incredibly essential, as some quantity of the separation energy is used within the receptor to bring a conformational amendment. This results in altered behavior of associate degree associated particle channel or the protein [13].

## II. LITERATURE SURVEY

Many research works are carried out with the complexes for which binding affinity is known. The purpose has been studied and the need is identified from literature survey. Xueling Li et al., proposed a method for automatic protein-protein affinity binding based on svr-ensemble. Two-layer support vector regression (TLSVR) model is used to implicitly capture binding contributions that are hard to explicitly model. The TLSVR circumvents both the descriptor compatibility problem and the need for problematic modeling assumptions. Input features for TLSVR in first layer are scores of 2209 interacting atom pairs within each distance bin. The base SVRs are combined by the second layer to infer the final affinities. This model obtains a good result [14].

Volkan Uslan, Huseyin Seker projected a technique for the quantitative prediction of HLA-B*2705 amide binding affinities mistreatment Support Vector Regression to achieve insights into its role for the Spondyloarthropathies. They proposed the prediction of human class I MHC allele HLA-B*2705 binding affinities using SVR, prior to processing, normalisation and feature selection were performed. The descriptors that form the amino acid composition were normalised to [0, 1] to ensure that every descriptor represented within the same range of values. Subsequently, Multi-Cluster Feature Selection (MCFS) method was used. the prediction of human class I MHC allele HLA-B*2705 binding affinities using SVR, prior to processing, normalisation and feature selection were performed. The descriptors that form the amino acid composition were normalised to [0, 1] to ensure that every descriptor represented within the same range of values. Subsequently, Multi-Cluster Feature Selection (MCFS) method was used [15].

From the background study it was perceived that most of the works were based on the complex for which binding affinity was provided with the database. This emphasizes the need of more research on affinity prediction with known structures and unknown drugs by designing and deriving the effective features for generating new model. Hence it is proposed to build a predictive model from which affinity can be predicted for docked complex. The study is carried out using 17 structures for different types of SCA and 18 ligands were used. Each structure is docked with ligand to prepare a complex and the features like sequence descriptors, rf-score, cyscore, energy based complexes, autodock vina scores are extracted to get the accurate recognition rate of affinity. The experiment is executed using learning models to predict affinity. The models are built with linear regression, polynomial regression, support vector regression, neural network regression and ridge regression coded in python using scikit learn framework.

## III. MATERIALS AND METHODS

The key idea in this research is to determine the rate of affinity from mutated structure based on repeats variants and to provide an effective regression models for affinity prediction. Structures are taken from pdb and mutation is inserted with the information available. Sequence based and structure based discriminative features are identified and it is used to train the model.

In this research work, the structures are synthesized based on the mutation position and its location given in HGMD database. For instance, in the database the mutational information for repeats is specified as 46-100 in the coding region. The structure is altered based on the information for all types of SCA structures specified in this paper.

Mutations will have many effects on the behaviour and copy of a macromolecule betting on wherever the mutation happens within the aminoalkanoic acid sequence of the macromolecule. If the mutation occurs in the region of the gene that is responsible for coding for the protein, the amino acid may be altered. When changes occur in amino acid coding region, formation of protein gets altered. Repeat mutation expands in the sequence and changes the structure of protein. Mutation in structure leads to various disorders. Muthukamarasamy et al., [16] analyzed homopeptide repeats on *Mycobacterium tuberculosis* H37Rv. In this sudy they analyzed how homopeptide repeats influences the protein structure and function.

### Docking

Docking is performed by preparing receptor (protein) and ligand (drug). Protein is manufactured by converting the protein into pdbq format that is by adding hydrogen, computing geastier charge and kolman charges are added. Protein.pdbq is converted to pdbqt by not adding partial charges to the protein because the protein possesses the charge. Protein is covered by grid box and size, centre of the box are noted and it is saved as grid parameter file. Ligand is prepared by converting the ligand into pdbqt format that is by detecting root in torsion tree. In search parameters genetic algorithm is used. Genetic algorithm uses global search, GA begin with a population of random ligand conformations in random orientations and at random translations. The quantity of people is within the population, will be able to try this victimisation “ ga_pop_size “: generally, fifty is found to be an honest worth. AutoDock counts variety|the amount|the quantity} of energy evaluations and also the number of generations because the moorage run proceeds: the run terminates if either limit is reached (” ga_num_evals “ and “ ga_num_generations “ respectively). You ought to critically set the quantity of the most effective people within the current population that mechanically survive into ensuing generation, victimisation “ ga_elitism “: generally this is often one. you ought to conjointly specify a correct worth for the rate of gene mutation using “ ga_mutation_rate “ and the rate of gene crossover “ ga_crossover_rate “; typically these are 0.02 and 0.80 respectively, although setting “ ga_crossover_rate “ to 0.00 reduces the genetic algortihm (GA) to an evolutionary programming (EP) method [17]. Each run will give one docked conformation. Output is saved as docking parameter file. The next step is to run autogrid which will generate grid log file and autodock generates docking log file. The docking log file is essential to check the docking results. The docking conformation is chosen based on which conformation is high and provide better result. For instance the 3D structure of ataxin-3, structure id is 2dos and the ligand interacted is bromocriptine and the docking conformation is shown in Fig 1. The amino acids that are interacted with ligand bromocriptine are ARG 124, GLN 110 and GLN 24.

**Fig 1.**
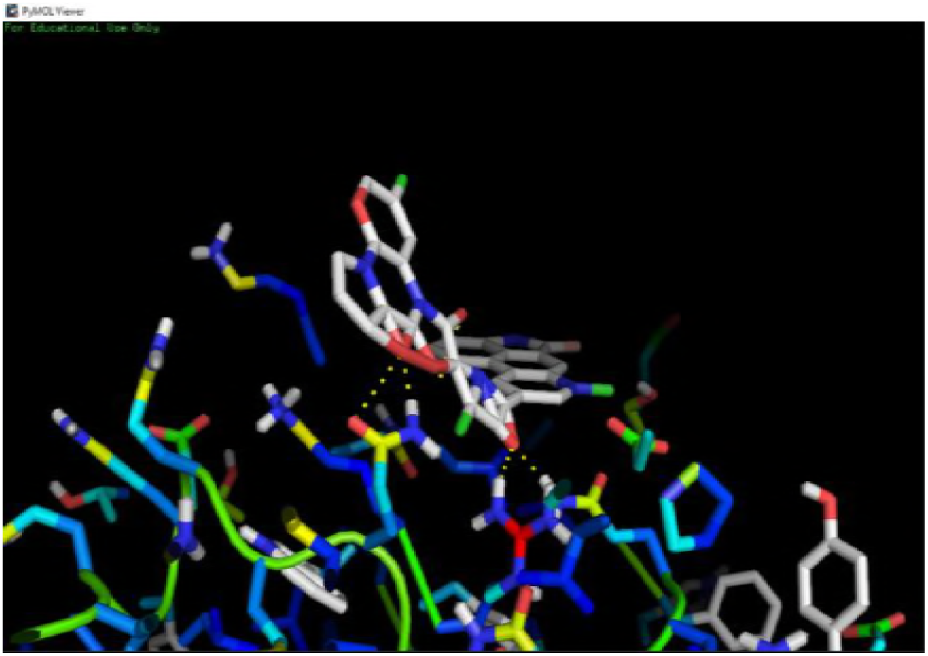
Docking Conformation.

### Explanatory variables and Response variables investigated in this study

The explanatory variable and response variables aids in improving the recognition rate. These variables are derived from structure and sequence based. Features that are examined in this analysis are classified into 5 categories. (1) Energy based (2) rf-score (3) Cyscore (4) Sequence descriptors and (5) Autodock vina scores.

#### Energy based descriptors

Energy based features aids in recoginizing the rate of affinity in a accurate manner. In [18] the author describes the work with rf-score features, which is not sufficient to predict the affinity. Hence features like binding energy, inhibition constant, intermolecular energy, vanderwaal’s hydrophobic desolvation energy, electrostatic energy, total internal energy and torsional energy. These energy based features are calculated using autodock tool. Energy based descriptors and its description are given in Table 2.

**Table 2.** Energy based descriptors and its Description.

| Features | Description |
| --- | --- |
| Binding Energy | Affinity of ligand-protein complex |
| Inhibition constant | Confidence of inhibitor |
| Intermolecular Energy | Energy between non-bounded atoms |
| Desolvation Energy | Energy lose of the interaction between substance |
| Electrostatic Energy | Amendment on the electricity non delimited energy of substance |
| Total Internal Energy | Change of all energetic terms |
| Torsional Energy | Dihedral term of internal |
|  | energy |

#### Rf-score

Rf-score has 36 features, along with energy based features these rf-score features aids in predicting the binding affinity. Rf-score features are extracted using python scripts. In [19] the author has determined the importance of rf-score for affinity prediction. In this work, each feature will comprise the number of occurrences of a particular protein-ligand atom type pair interacting within a certain distance range. Our main criterion for the selection of atom types was to generate features that are as dense as possible, while considering all the heavy atoms that are commonly observed in PDB complexes. As the number of protein-ligand contacts is constant for a particular complex, the more atom types are considered the more sparse the resulting features will be. Therefore, a minimal set of atom types was selected by considering atomic number only. Furthermore, a smaller set of intermolecular features has the additional advantage of leading to computationally faster scoring functions.

Here we consider nine common elemental atom types for both the protein P and the ligand L:

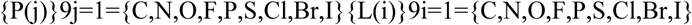

The occurrence count for a particular j-i atom type pair is evaluated as:

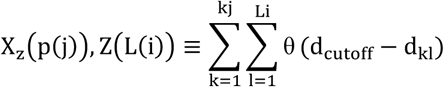

where d_kl_ is the Euclidean distance between k^th^ protein atom of type j and the l^th^ ligand atom of type i calculated from the PDBbind structure; K_j_ is the total number of protein atoms of type j and L_i_ is the total number of ligand atoms of type i in the considered complex; Z is a function that returns the atomic number of an element and it is used to rename the feature with a mnemonic denomination; Θ is the Heaviside step function that counts contacts within a d_cutoff_=12Å neighbourhood of the given ligand atom. For example, x_7,8_ is the number of occurrences of protein nitrogen interacting with a ligand oxygen within a 12Å neighbourhood. This cutoff distance was suggested in [19] as sufficiently large to implicitly capture solvation effects, although no claim about the optimality of this choice is made. This representation leads to a total of 81 features, of which 45 are necessarily zero across PDBbind complexes due to the lack of proteinogenic amino acids with F, P, Cl, Br and I atoms. Therefore, each complex will be characterised by a vector with 36 features:

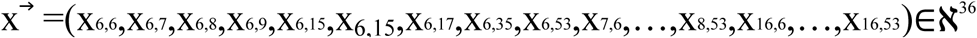

#### Cyscore

Cyscore is an empirical scoring function consists of four numerical features. In [20] the author compared the cyscore scoring functions with other scoring functions, in which cyscore gives the better predicition. Hence features like Cyscore, hydrophobic energy, van der Waals interaction energy, hydrogen-bond interaction energy and the ligand’s conformational entropy are captured from the complexes using pyhton script. Cyscore descriptors and its description are depicted in Table 3.

**Table 3.** Cyscore and its description.

| Features | Description |
| --- | --- |
| Hydrophobic free energy | Driving force for folding of globular proteins |
| Cyscore | Scoring function |
| Van der waals interaction energy | Electrical interactions between two or more atoms |
| Hydrogen-bond interaction | Bond is shared between two electronegative atoms |
| Ligand's conformational | Measures change in free |
| entropy | energy upon binding |

#### Sequence Descriptors

Sequence Descriptors have many features which aids in identifying the affinity. In [21] the author used atom pair occurrence and distance-dependent atom pair features with PLS technique. In this work the commonly used descriptors are captured with the package protr in R and coded in R. Commonly used descriptors are amino acid composition, autocorrelation, CTD, Conjoint Triad, Quasi-sequence-order descriptors, Pseudo amino acid composition (PseAAC), Profile-based descriptors. Sequence descriptors and its description are explained in Table 4.

**Table 4.**
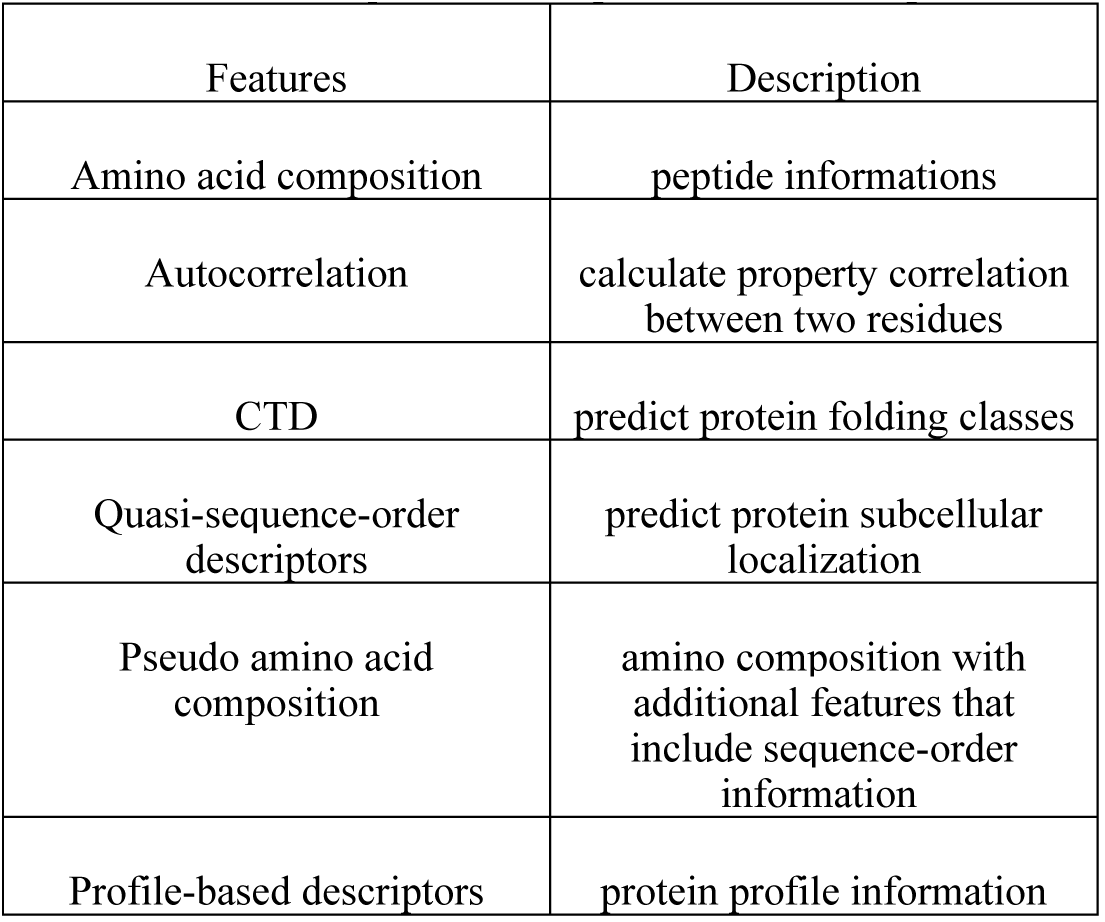
Sequence Descriptors and its description.

#### Autodock Vina scores

Autodock vina scores is a hybrid scoring function combination of empirical and knowledge based scoring function. In docking it improves the accuracy and speed with scores, efficient optimization and multithreading [21]. Hence features like binding affinity, gauss 1, gauss2, repulsion, hydrophobic and hydrogen bonding are considered and defined with the command line. Autodock vina scores and its description are given in Table 5.

**Table 5.** Autodock Vina scores and its description.

| Features | Description |
| --- | --- |
| $\Delta G_{\text{gauss}}$ | Attractive term for dispersion, two Gaussian functions |
| $\Delta G_{\text{repulsion}}$ | Square of the distance |
| $\Delta G_{\text{hydrophobic}}$ | Ramp function |
| $\Delta G_{\text{Hbond}}$ | Ramp function for interactions with metal ions |

## IV. PREDICTIVE ALGORITHMS

Affinity prediction models are built using various regression algorithms. In this work, models are built using linear regression, polynomial regression, ridge regression, support vector regression and neural network regression. These regression algorithms are coded with python in scikit-learn environment. The algorthims are discussed below in detail.

### 4.1 Linear Regression

It is a linear approach for modeling the connection between dependant variable y and one or additional informative variable x. If it’s one informative variable then it’s easy regression toward the mean, and once the informative variable is 2 or additional then it’s referred to as multiple regression toward the mean. The equation for linear regression is

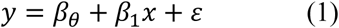

Linear regression is about finding best fitting straight line through the points. The best fitting line is called regression line. The regression line for my data is given in fig 2. Since the mean squared error is less the regression line is fitting thorugh the points.

**Fig 2.**
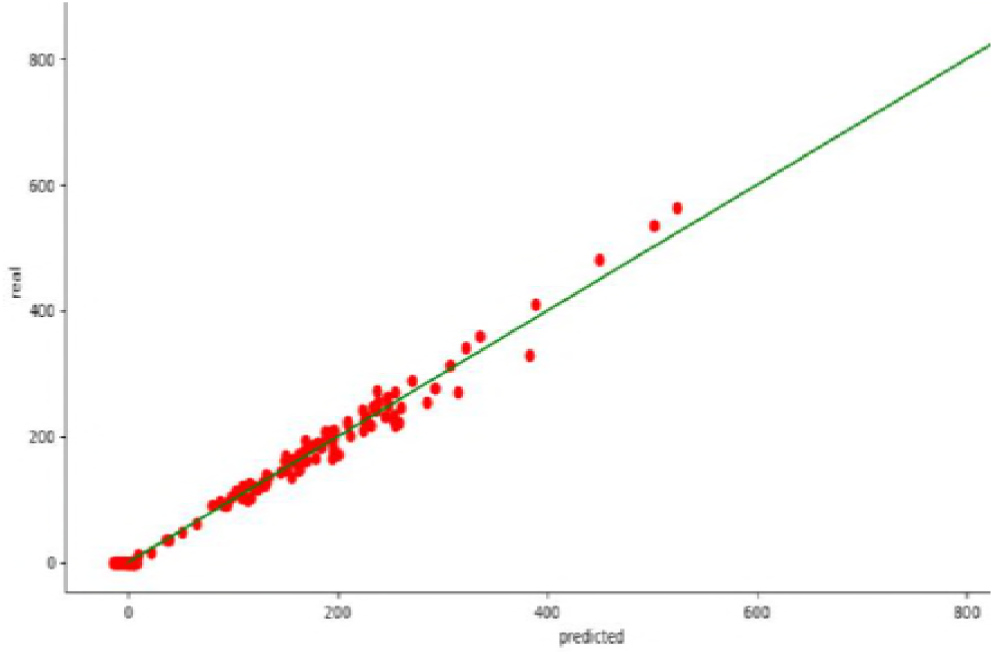
Regression line of linear model.

### 4.2 Polynomial Regression

Polynomial regression is special case of linear regression. It fits the non-linear relationship between the independent variable and conditional mean of dependent variable. Polynomial regression adds the cubic or quadertic form to the linear regression. In polynomial regression the expected model of y as nth degree polynomial, yielding the general polynomial regression model

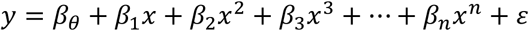

A cubic polynomial regression fit for the dataset is given in Fig 3.

**Fig 3.**
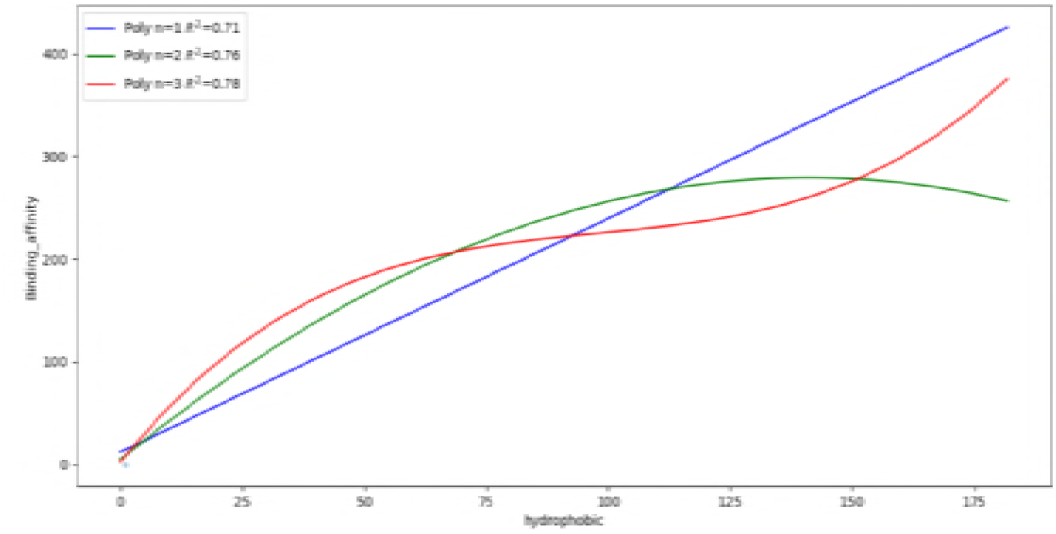
Cubic Polynomial fit.

### 4.3 Ridge Regression

Ridge Regression could be a technique for analyzing multiple correlation information that suffer from multiple regression. once multiple regression happens, method of least squares estimates area unit unbiased, however their variances area unit massive so that they could also be aloof from verity price. By adding a degree of bias to the regression estimates, ridge regression reduces the quality errors. It performs L2 regularization reduction objective = LS Obj + α * (sum of sq. of coefficients). reduction objective = LS Obj + α * (sum of sq. of coefficients).

Here, α (alpha) is the parameter which balances the amount of emphasis given to minimizing RSS vs minimizing sum of square of coefficients. α can take various values. The figure for ridge regression with the function of regularazation is given in fig 4.

**Fig 4.**
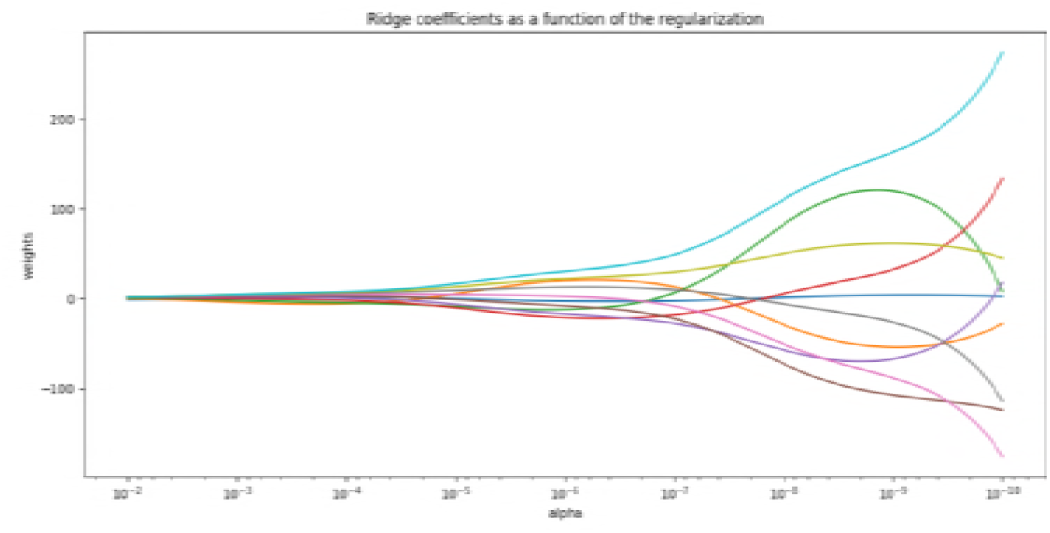
Ridge regression.

### 4.4 Support Vector Regression

Support vector machines tackle classification and regression issues of non-linear mapping computer file into high dimensional feature areas, whereby a linear call surface is intended. Support vector regression algorithms area unit supported the results of the applied mathematics theory of learning given by vapnik, that introduces regression on the fitting of a record v to the information. SVM regression estimates the worth of w to get the function.

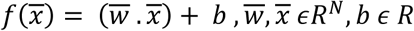

By introducing ε insensitive loss function as

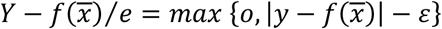

It retains all the properties that characterize maximal margin algorithms of support vector machines such as duality, sparseness, kernel and convexity. It has become a powerful technique for predictive data analysis with many applications in varied areas of study like biological contexts, drug discovery, civil engineering, sunspot frequency prediction, image tracking, image compression etc.,

The model made by support vector regression (SVR) solely depends on a set of the coaching information, as a result of the price operate for building the model ignores any coaching information that square measure near the model prediction.

The support vector regression uses a price operate to live the empirical risk so as to reduce the regression error. though there square measure several selections of the loss functions to calculate the price, e.g., least modulus loss operate, quadratic loss operate, etc., the ε inability loss operate is such a operate that exhibits the sparsely of the answer [22].

### 4.5 Neural Network Regression

A multilayer perceptron (MLP) may be a category of feedforward artificial neural network. associate MLP consists of a minimum of 3 layers of nodes. apart from the input nodes, every node may be a nerve cell that uses a nonlinear activation operate. MLP utilizes a supervised learning technique known as backpropagation for coaching.

#### Activation function

If a multilayer perceptron contains a linear activation operate all told neurons, that is, a linear operate that maps the weighted inputs to the output of every vegetative cell, then algebra shows that any variety of layers is reduced to a two-layer input-output model.

The 2 common activation functions ar each sigmoids, and ar delineated byThe initial could be a hyperbolic tangent that ranges from −1 to one, whereas the opposite is that the supply operate,which is analogous in form however ranges from zero to one.

The output of the ordinal node (neuron) and is that the weighted total of the input connections. various activation functions are projected, together with the rectifier and softplus functions. additional specialised activation functions embody radial basis functions (used in radial basis networks, another category of supervised neural network models).

#### Layers

The MLP consists of three or more layers (an input and an output layer with one or more hidden layers) of nonlinearly-activating nodes making it a deep neural network. Since MLPs are fully connected, each node in one layer connects with a certain weight to every node in the following layer.

#### Learning

Learning occurs in the perceptron by changing connection weights after each piece of data is processed, based on the amount of error in the output compared to the expected result.

Scoring parameters for the regression are evaluated using 10-fold cross validation using metric functions of scikit library which is tabluated in Table 5 and in fig 5. Support Vector regression gives the highest variance score.

**Fig 5.**
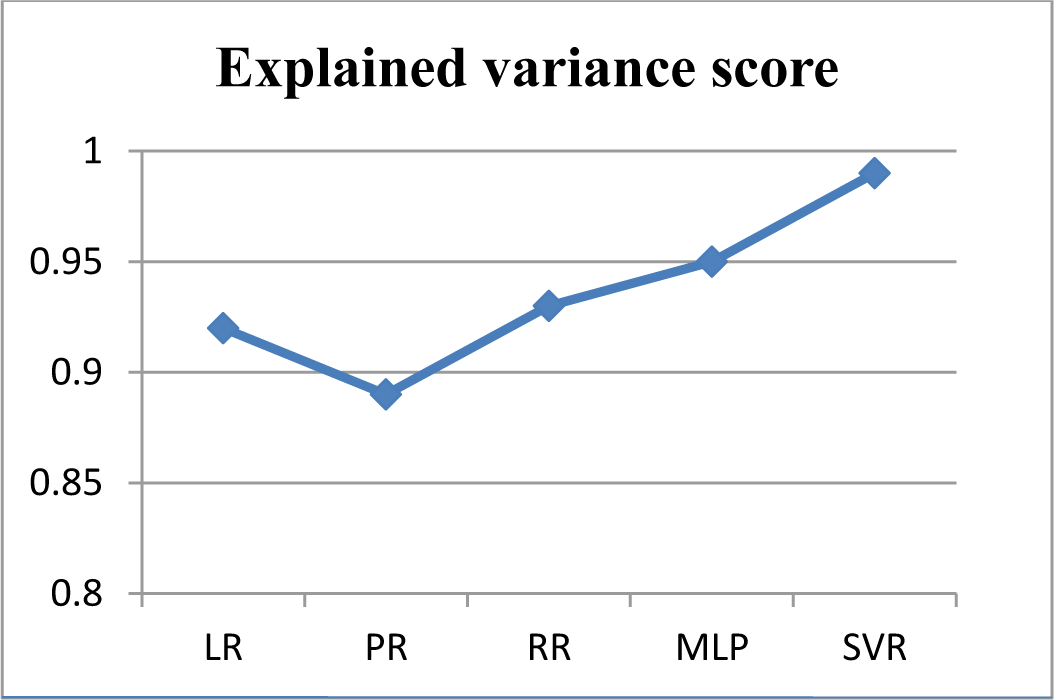
Variance score of learning models.

## V. Experiment and Results

An efficient affinity prediction models are built using regression techniques in scikit learn environment. The scikit learn framework uses python library for implementing the regression problem [23]. The structures are mutated based on the mutation information. Discriminative features were identified and captured from the docked complex as described in section 2. The training data set with instances related to six categories of spinocerebellar ataxia that is ataxin-1, ataxin-2, ataxin-3, ataxin-6, ataxin-8 and ataxin-10 has been developed. The volume of the dataset accounts to 306 complexes from the docked complex of 17 structure and 18 lignads to work with the statiscal learning techniques. All six categories have more than one protein structure.

The main virtue of statiscal machine learning methods is their ability to model high dimensional datasets which are used increasingly for data analysis. Scikit-learn contains an extremely hefty set of statistical learning algorithms, for both supervised and unsupervised learning. The scikit-learn tool kit has a variety of algorithms, that is matched by few packages, and it is implemented in python, that matched perfectly in the rich python ecosystem. The scientific libraries that are used for machine learning are numpy, scipy, matplotlib. Scikit-learn accept all objects and algorithms in the size of 2-dimensional arrays to make the data as generic and domain-independent. Depending on the purpose scikit-learn objects constitute the uniform set of methods such as estimators, predictors and transformers. In anaconda framework, the conda comprises of various library packages that are required for implementation. The extracted descriptors from the complexes are stored in .csv files. The .csv files are converted into a numpy array, as scikit-learn library accepts a numpy array in its implementation. The data frame is built with the numpy array. Normalizing the data by transforming the feature values into the range between 0 and 1 aid in scaling the input attributes for a model.

Predictive models based on Linear Regression, Polynomial Regression, Ridge regression, Support Vector Regression and Neural Network Regression are developed using python library framework in scikit learn. While learning SVR model, the cost, gamma and kernel parameters are tuned to obtain good results. The predictive results of various learning models are shown in Table 6 and Figures 2-10.

**Table 6.** Perfomance results of Regression models.

| Regression models | Explained variance score | Mean squared error | R2_score | Mean absolute error | Median absolute error | Mean squared logarithmic error |
| --- | --- | --- | --- | --- | --- | --- |
| Linear Regression | 0.92 | 14.58 | 0.85 | 0.7 | 0.7 | 0.059 |
| Polynomial Regression | 0.89 | 18.21 | 0.78 | 0.78 | 0.65 | 0.056 |
| Ridge Regression | 0.93 | 13.45 | 0.86 | 0.8 | 0.7 | 0.048 |
| MLP Regressor | 0.95 | 15.05 | 0.88 | 0.75 | 0.6 | 0.044 |
| SVR Regression | 0.99 | 9.05 | 0.90 | 0.6 | 0.5 | 0.039 |

**Fig 6.**
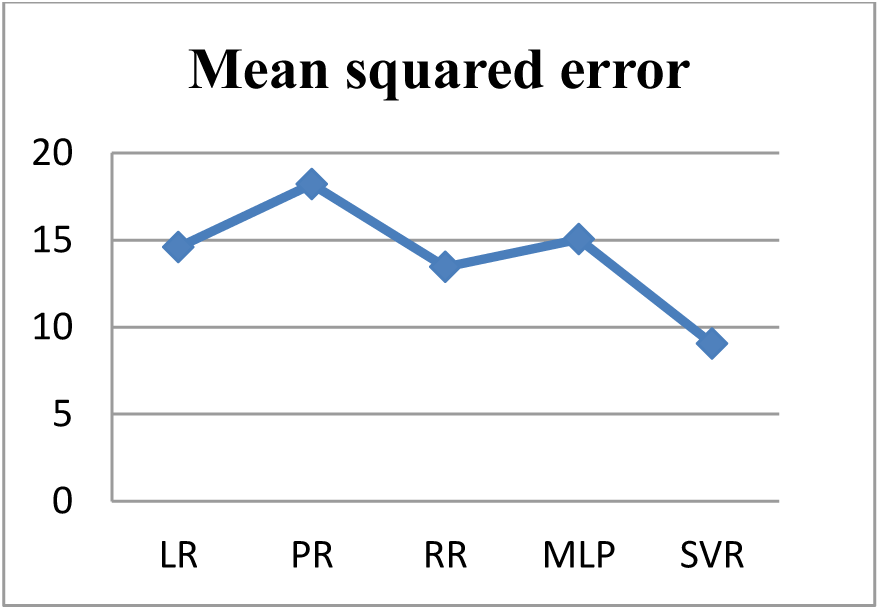
Mean squared error of learning models.

**Fig 7.**
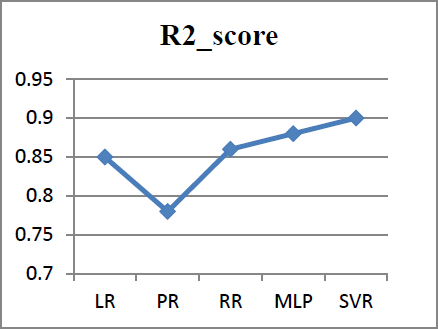
R2_score of learning models.

**Fig 8.**
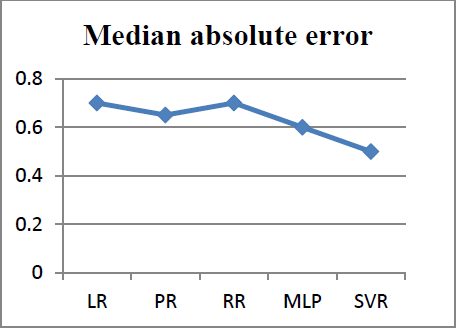
Median absolute error of learning models.

**Fig 9.**
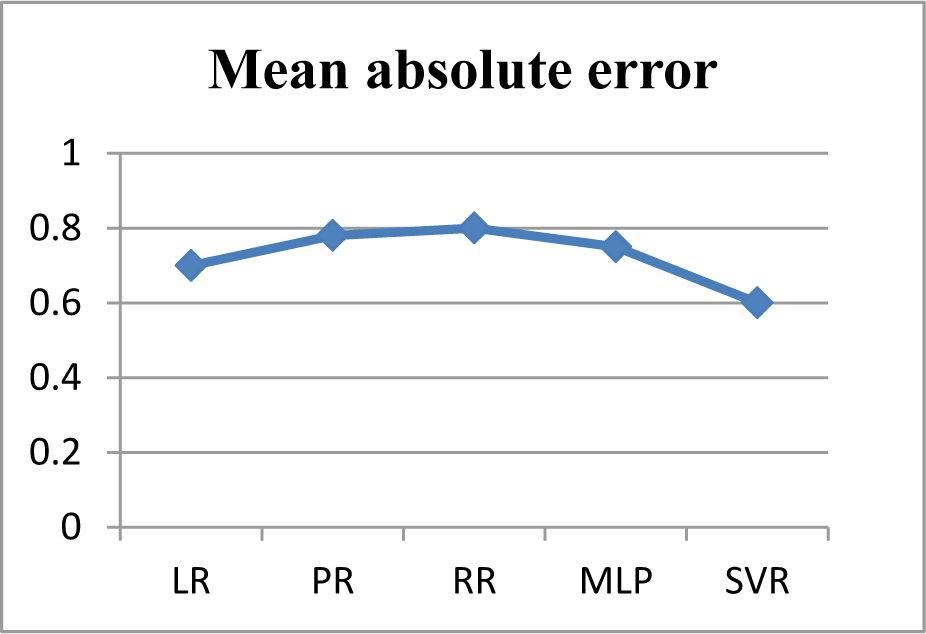
Mean absolute error of learning models.

**Fig 10.**
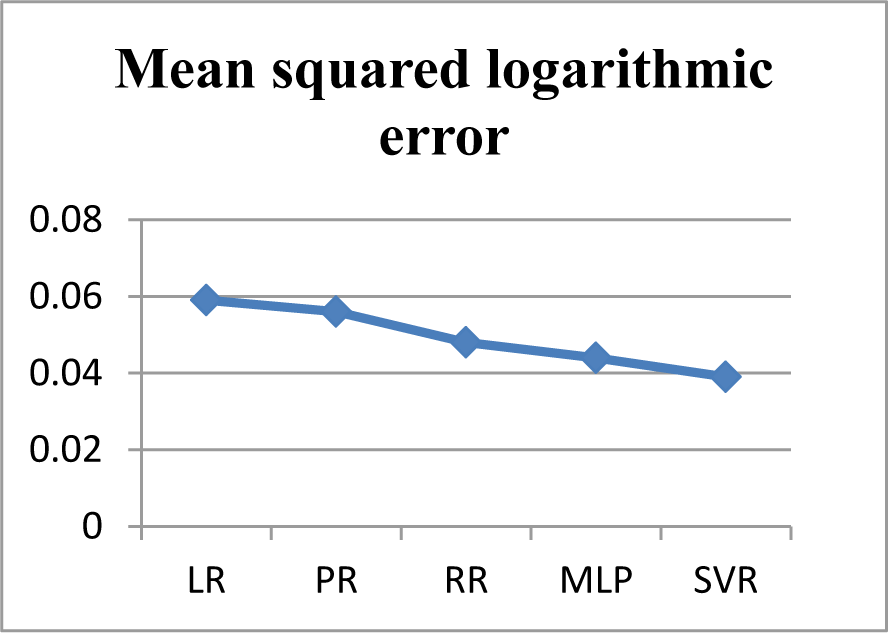
Mean squared logarithmic error of learning models.

From the above results, it was observed that Support Vector Regression performs well with the minimum error rate and with high variance score. The regression metrics are depicted in the figures from 5-10. The figures clearly shows that the error rate achieved by SVR is comparatively low, when compared to other predictive algorithms. Among all the metrics, explained variance score should be highest to measure the strength of prediction and Support Vector Regression gives the explained variance score as Mean squared error should be very minimum, in order to achieve better model. Support vector regression gives the minimum error rate of 9.05. The other regression metrics like R2_score is also high in support vector regression. Error metrics like median absolute error, mean absolute error and mean squared logarithmic error such as 0.5, 0.6 and 0.039 respectively are obtained by SVR which is comparatively low, when compared to other predictive algorithms like LR, PR, RR and MLP.

In this work, many predictive measures are analyzed like median absolute error, mean squared logarithmic error, mean absolute error and R2-score in addition with mean squared error and explained variance error. From the above inference, it proves support vector regression performs high with the better model and with minimal error rate.

## VI. CONCLUSION

Binding Affinity prediction task is a regression problem to identify the affinity from mutated protein structures. Energy-based descriptors, autodock vina scores, cyscore and rf-score are proposed and extracted from the complexes. An eminent model is built using statistical machine learning technique through scikit learn in anaconda framework. The performance of the predictive algorithms are increased by employing different estimators from the python library. The predictive algorithms were evaluated based on various metrics and the results indicate that the support vector regression implemented in this framework is best suited for predicting the binding affinity. Repeat mutation is also considered to build models through statistical learning technique and their results are analyzed. This work proves that the learned model is highly efficient in the prediction of binding affinity.

